# BIOCOMPATIBILITY OF LARGE-AREA 2-DIMENSIONAL ELECTRONIC MATERIALS WITH NEURAL STEM CELLS

**DOI:** 10.1101/2025.07.19.665698

**Authors:** RT Jayanth, Rebecca Duquette, Shanmukh Kutagulla, Sabrina Pietrosemoli Salazar, Emmanuel Okogbue, Jingyuan Zhou, Yeonwoong Jung, Xiangfeng Duan, Dmitry Kireev, Stephanie K. Seidlits, Deji Akinwande

## Abstract

Two-dimensional (2D) electronic materials hold immense promise for next-generation bio/neuro-electronic interfaces, but their biocompatibility has remained uncertain due to conflicting reports from studies focused on exfoliated flakes and suspensions. In this work, we present a comprehensive *in vitro* evaluation of electronic-grade large-area, chemical vapor deposition (CVD)-grown 2D materials – including platinum diselenide (PtSe_2_), platinum ditelluride (PtTe_2_), molybdenum disulfide (MoS_2_), and graphene – as substrates for mouse neural stem cell culture. Across all CVD-grown materials, the stem cells exhibited outstanding viability, with no significant differences in metabolic activity or live/apoptotic cell ratios compared to laminin-coated glass controls (p > 0.05). Importantly, these large-area 2D materials robustly supported neuronal differentiation, as evidenced by widespread βIII-tubulin expression. Strikingly, we found that flaky MoS₂ promoted significantly greater neuronal maturation (>75% NeuN⁺ neurons) than any other substrate tested (25–50% NeuN⁺; p < 0.05), revealing the critical influence of material format on bioactivity. While PtSe₂ showed a tendency to promote glial lineage differentiation, our findings firmly establish large-area CVD-grown 2D materials as biocompatible, tunable platforms for neural interfacing, paving the way for their integration into advanced bio/neuro-electronic devices.

## 1 INTRODUCTION

Bioelectronic materials for long-term, *in situ* measurements should be mechanically flexible, cause minimal cell or tissue damage and enable robust electrophysiological measurements. However, traditionally used metals and metal-alloys, including platinum, platinum-iridium, gold, titanium, tungsten, and stainless steel have low mechanical flexibility and show increased cytotoxicity with longitudinal usage. While these materials can support glial cell growth and promote neuronal morphological changes, they suffer from decreased signal-to-noise (SNR)^1,2,3,4^. For rodent and non-human-primate neural electrophysiology, silicon probes and metal-alloy electrodes are extensively used in research. However, these electrodes are especially vulnerable to degradation over time, compromising the long-term recording performance and the health of the animal^5^. As these electrodes are usually rigid and have lower surface-to-volume ratio, they are prone to causing tissue damage and cannot be used longitudinally to probe deeper regions of the brain. Hence, mechanical flexibility, longitudinal biocompatibility, and robust electrical, thermal, and electromagnetic-wave conductivity are crucial requirements for future biological and bio-electronic materials.

In all these facets, two-dimensional (2D) electronic materials have shown remarkable characteristics. They are highly mechanically flexible, with tunable electronic, thermal, and electromagnetic-wave conductivity profiles. However, there is typically a trade-off between physical performance of any material (2D or otherwise) and its biocompatibility with interfaced cells. In most instances, these materials are stacked in layers adhered with strong in-plane covalent bonds and weak out-of-plane van der Waals bonds^6^. Since the discovery and characterization of graphene, a 2D atomic-sheet of carbon atoms, there has been widespread basic and applied scientific research to understand its physical attributes^7^. Graphene has been a widely used as an *in vitro* substrate for cell culture and has been shown to enhance synaptic activities of primary neurons^8,9^ and mesenchymal stem cells (MSCs) compared to standard tissue culture plastic (TCP) and glass^10,11^. Large-area chemical vapor deposition (CVD)-grown graphene has also been incorporated as a sensing material for *in vivo* neural electrodes^12^. However, conflicting evidence exists on how graphene might affect inherent neuronal electrophysiological properties (such as affecting the excitatory activity of pyramidal cells), and the overall long-term neural-network deficits with exposure to graphene^12,13^ (see Table S2). Another disadvantage lies in the fact that graphene does not have a direct band-gap in its intrinsic state which makes it harder to use as a high SNR bioelectronic material^7^. One might have to add dopants to graphene to make it semi-conducting but that might compromise the mechanical and thermal properties of its pristine version^14^. Hence, there is a need to explore alternative 2D materials that have a wide range of electronic, mechanical, and thermal properties^15^.

Beyond graphene, transition-metal-dichalcogenides (TMDs), which have a generic MX_2_ composition (M – Metal, X – Chalcogen), have emerged as an exciting class of 2D materials^16^. Most TMDs are either semiconducting, metallic, insulating, or superconducting and exhibit topological electronic phases^5^. MoS_2_ is the most extensively studied 2D TMD owing to its more straightforward growth process using CVD^17^. Electronically, its CVD-grown form is known to be an indirect band gap semiconductor, whereas its monolayer/few-layer form has a direct band gap^18^. This thickness-dependent electronic property of MoS_2_, along with its mechanical flexibility, has made it useful for flexible bioelectronic devices^19^. Similarly, PtTe_2_ and PtSe_2_ (Pt-TMDs in general) have thickness-dependent semiconductor-metal transition properties and have excellent air- and water-stability compared to other TMDs^20,21^. These properties position Pt-TMDs as prime candidates for future flexible bioelectronic devices^22^.

Biocompatibility of TMDs typically has been assessed based on exfoliated forms of the material in liquid suspensions, rather than their large-area 2D surfaces^23–26^. A common observation across studies using suspension preparations of exfoliated TMD flakes is material cytotoxicity above a critical concentration, with additional dependencies on flake size and thickness^23^. It remains unclear whether such exfoliated flakes possess sufficient surface contact-area for cell adherence and viability. It is also possible that the flakes can be readily internalized by cells, leading to cell death^25^.

Although large-area, electronics-grade graphene and MoS_2_ have been grown previously using CVD, this format has only recently been achieved with PtSe_2_ and PtTe_2_^27^. Therefore, we designed this study to directly compare the bioelectronic interfacing potential of large-area 2D materials (graphene, PtTe_2_, PtSe_2_, MoS_2_), flaky form of MoS_2_ (hereafter named flaky MoS_2_), and thin layers of metals that include gold and platinum, using embryonic spinal cord-derived mouse neural stem cells (mNSCs) (Sch 1). All substrate materials support mNSC viability at least as well as the laminin-coated glass control. Morphological differences of the adhered cells were observed for CVD-grown MoS_2_ and PtTe_2_ substrates, on which cells showed less overall area coverage and spreading. Compared to other conditions including glass substrate controls, mNSCs on flaky MoS_2_ differentiated into more neurons, while those on PtSe_2_ tended to become glial cells, including oligodendrocyte-like cells and astrocytes. This study provides a comprehensive view of the potential of 2D materials with appropriate electronic properties, mechanical strength and flexibility to be utilized for bioelectronic, neural interfaces that can inform the choice of materials for future bioelectrode designs.

**Scheme 1.**
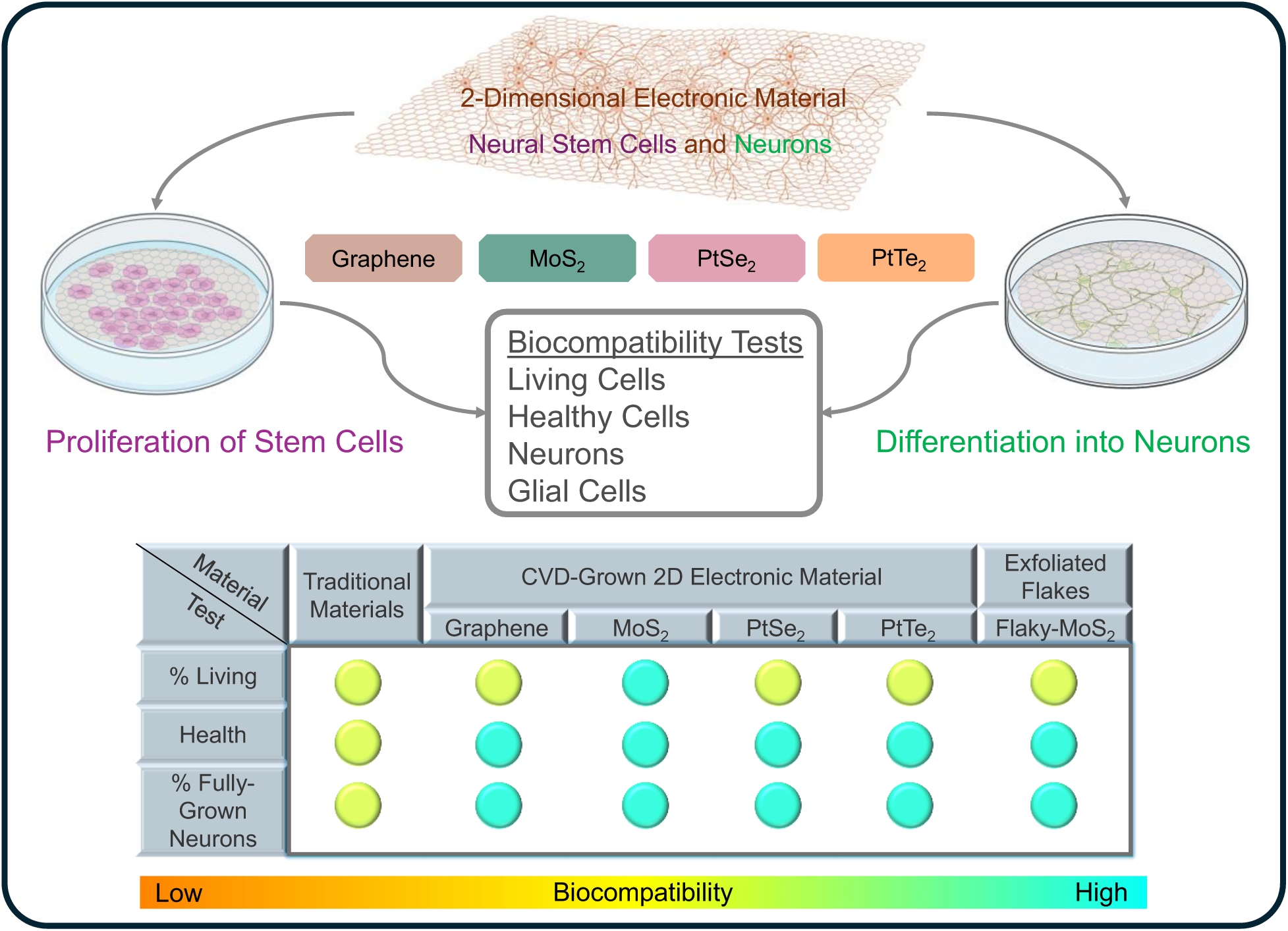
Schematic illustration of the experimental set-up and the main results of the biocompatibility studies.

## 2 MATERIALS AND METHODS

All the materials of interest (graphene, PtSe_2_, PtTe_2_, MoS_2_, and flaky MoS_2_) used in this study had glass as the substrate.

### 2.1 Growth and Transfer of Graphene

Single-layer high-quality CVD grown graphene was purchased from Grolltex. The received graphene/Cu foil was placed over a SiO_2_/Si wafer ensuring that the edges were properly sealed using a polyimide (PI) tape. A thin layer of polymethyl methacrylate (PMMA, 950 A4) was spin-coated at approximately 2500 rpm for 60 seconds eventually forming a thickness of 200 nm. This assembly was annealed at a temperature of 150°C for 5 minutes and the PI tape was carefully removed. The copper foil was placed into the etchant solution (0.1M (NH_4_)_2_S_2_O_8_) for a duration of at least 8-12 hours. After Cu was completely etched, the PMMA/graphene film was transferred into deionized (DI) water to be cleaned thoroughly. Clean glass slips (9 mm diameter) were rinsed with DI water and air-dried prior to the transfer of PMMA/graphene stack. Once dried, the stack was annealed for 5 minutes at 150°C to help the re-flow of PMMA ensuring better graphene-substrate adhesion. Lastly, the PMMA layer was removed from the graphene with an acetone bath and rinsed in an isopropyl alcohol (IPA) bath. The film was then dried gently using a nitrogen gun, placed and glued atop a glass coverslip using Polydimethylsiloxane (PDMS, Avantor) prior to mounting into a 48-well plate for cell culture.

### 2.2 Growth and Transfer of MoS_2_

MoS_2_ was grown using a 2-step sulfurization process directly onto glass slips^17^. Glass slips were first immersed in acetone for 1 hour and rinsed with isopropanol (IPA) for cleaning. Glass slips were adhered to water soluble tape in batches of ∼20. Molybdenum metal (0.7 nm) was first evaporated onto the glass slip/water soluble tape stack in a cleanroom (CHA Industries). Immediately following, the stack was immersed in DI water to dissolve the tape. To further remove contaminants, the slips were individually rinsed twice with DI water. These slips were then loaded into an alumina crucible (AdValue Tech) and loaded into the middle zone (Zone 2) of a 3-zone tube furnace. Upstream in zone 1, a crucible containing 3.5 g of sulfur powder (Sigma Aldrich) was loaded, and the tube sealed. 125 sccm of argon was then flowed, and the tube was purged 3 times to remove oxygen and moisture, until the pressure of the tube settled at ∼50 mTorr. Then the sulfur zone (Zone 1) was heated to 220°C, and Zone 2 was heated to 450°C. These conditions were held for 15 minutes to allow for MoS_2_ growth, and the argon flow was stopped and the tube allowed to come to room temperature. These samples were then unloaded for further characterization.

### 2.3 Flaky MoS_2_

A MoS_2_/IPA ink solution was prepared through a molecular intercalation/exfoliation method^28^. This process utilized a two-electrode electrochemical cell comprising a thin cleaved MoS_2_ crystal (molybdenite) as the cathode, a graphite rod as the anode, and tetraethylammonium bromide (THAB, 98% from Tokyo Chemical Industry (TCI) dissolved in acetonitrile (40 ml; 5 mg/ml or higher) as the electrolyte. In the intercalation step, a negative voltage (5-10 V) was applied to the MoS_2_ cathode for 1 hour, facilitating the insertion of positively charged THA+ ions into the crystal and resulting in a fluffy CVD-grown material. This material was then rinsed with absolute ethanol and subjected to sonication in a 40 ml 0.2 M polyvinylpyrrolidone/dimethylformamide (PVP/DMF) solution (PVP: molecular weight approximately 40,000, Sigma-Aldrich) for 30 minutes to achieve a greenish nanosheet dispersion. Subsequently, the dispersion underwent centrifugation and was washed with IPA twice to eliminate excess PVP. Acetonitrile was used in the 1st step for initial intercalation of THAB. Following the steps to fully disperse MoS_2_ flakes, including 1 step of sonication in DMF and 3 steps of solvent exchange to remove unnecessary particles (organic and inorganic), the final product is MoS_2_ flakes dispersed in IPA. Then, the flaky MoS_2_ suspension was placed atop glass slips are spin-coated at 1000 RPM and annealed at a temperature of 150°C for 10 minutes.

### 2.4 Growth of Pt-TMDs over Glass

PtSe_2_ and PtTe_2_ layers were grown on SiO_2_/Si-Glass substrate using a two-step thermal assisted conversion process^27^. Electron beam deposition (Temescal) was used to deposit Pt films of desired thickness at a rate of 0.1 Å/s onto the glass substrates. The Pt-covered substrates were then placed in the middle zone of a CVD tube furnace with alumina boat containing tellurium (Te) or selenium (Se) placed at the upstream region for tellurization or selenization to obtain PtTe_2_ or PtSe_2_ layers, respectively. The CVD tube was pumped down to a low pressure of 25 mTorr followed by purging with Ar gas at a high flow rate to remove the moisture and residual gases. For both PtTe_2_ and PtSe_2_, the furnace was heated to the growth temperature of 400°C and held there for 50 min under a constant flow of Ar. The furnace was left to cool to room temperature naturally. These samples were then unloaded for further characterization.

### 2.5 Raman Spectroscopy

Raman spectroscopy was performed in a Renishaw inVia micro-Raman system. The excitation wavelength of 442 nm with an incident beam power of ∼1 mW and exposure time of 10 s was used for Raman measurement.

### 2.6 Water Contact Angle (WCA) Measurements

WCA measurements on 2D material over glass samples were carried out using a goniometer (Rame Hart 90-U3-PRO), and the same water volume was used to ensure uniformity across all measurements. Pre-culture samples were rinsed in IPA and DI water gently before using a nitrogen gun to dry out the samples. Samples underwent pre-culture WCA measurements and were then used as substrates for cell culture and analysis (methods 2.7 - 2.10).Afterwards, TrypLE Express (Gibco, 12604013) was added to the wells for 10 minutes at 37°C to detach the cells. Substrates were then rinsed three times with DI H_2_O and then one final time with 70% ethanol (EtOH). Post-culture substrates were dried and stored at room temperature before acquiring post-culture WCA measurements (n=5).

### 2.7 Preparation of 2D Electronic Materials for Cell Culture

2D material substrates on glass slips supports (circular substrate, 0.16 cm^2^ in area) were transferred onto the bottom of individual wells in a 48-well, tissue culture-treated well plate (Fig. 1, Fig. S4). Glass slips were adhered to the bottom of the well plates with the 2D electronic substrates facing upwards using polydimethylsiloxane (PDMS, Sylgard 184, 10:1 mix ratio). After PDMS curing (110°C for ∼20 minutes), the well plates were sterilized by placing them in an A2 biosafety cabinet under UV irradiation for 30 minutes, followed by incubation with 70% EtOH for 10 minutes. Next, EtOH was removed, and each substrate was rinsed twice with sterile Dulbecco’s Phosphate Buffered Saline (DPBS, Millipore Sigma, D8537).

**Figure 1.**
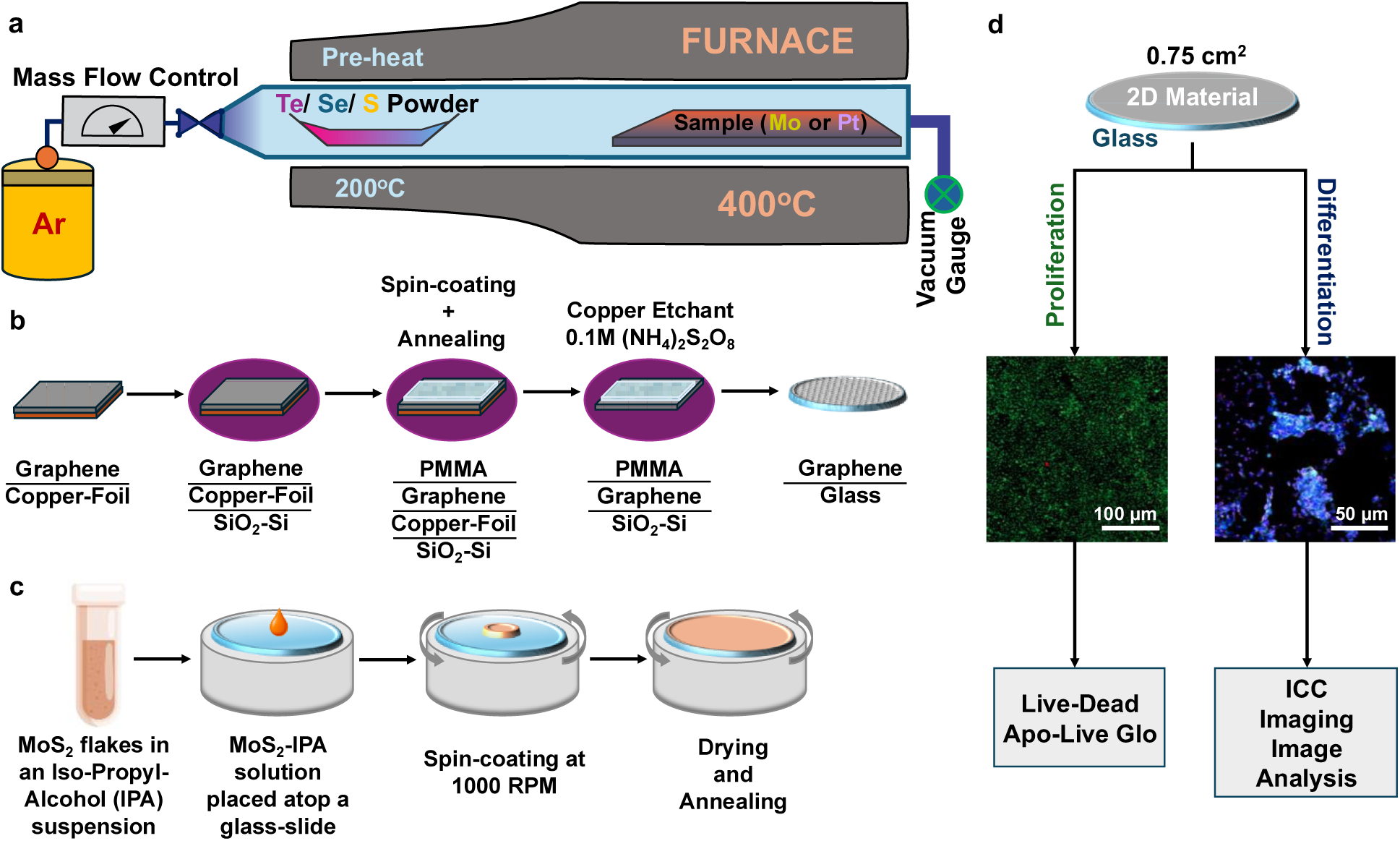
The preparation of samples and the experimental set-up. a) CVD growth procedure of PtSe2, PtTe2, and MoS2 is depicted in a schematic containing the CVD chamber, the growth temperatures, the substrate materials, and the precursors, b) process of transferring CVD-grown graphene layers onto glass coverslips, c) process of spin-coating and annealing the flaky MoS2-IPA suspension, d) flow-chart of the experimental set-up.

### 2.8 Cell Culture

Cryopreserved mouse spinal cord neural stem cells (mNSCs, E15-18, C57/Bl6) were purchased from Millipore Sigma (SCR031), expanded and used from passage 4-5 for all experiments. mNSCs were maintained in culture on poly-L-ornithine (PLO)/laminin-coated substrates in either proliferation medium (0.02 µg/mL epidermal growth factor (Thermofisher Scientific, AF10015100), 0.02 µg/mL basic fibroblast growth factor (Thermofisher Scientific, 100-18B-100UG), 2 µg/mL heparin (Millipore Sigma, H6279-25KU) and 0.2x (12µg/mL penicillin, 20 µg/mL streptomycin and 0.05 µg/mL Amphotericin B) antibiotic/antimycotic (ABAM) solution (Millipore Sigma, A5955-20mL) in NSC basal medium (Millipore Sigma, SCM003) or differentiation medium (1 µM retinoic acid, Millipore Sigma, R2625-50mg), 5 µM forskolin (Millipore Sigma, F3917-10mg), and 0.2x ABAM solution in NSC basal medium and maintained in an incubator (37°C, 5% CO_2_) with media changes every other day. Culture substrates were prepared by first coating clean material- or non-material-coated glass coverslips with PLO (Millipore Sigma, P4957-50 mL) (10 mg/mL in distilled, deionized water) and incubating for one hour at 37°C. Next, the coverslips were washed three times with sterile distilled, DI water. Then, a laminin (Millipore Sigma, CLS3544232-1EA) solution (6 µg/mL in DPBS) was added to coverslips for 1 hour at 37°C. For all studies, laminin was aspirated and mNSCs (700 cells/mm^2^) were immediately seeded onto PLO/laminin-coated 2D electronic material substrates (platinum, gold, graphene, PtTe_2_, PtSe_2_, MoS_2_, and flaky MoS_2_), PLO/laminin-coated coverslips, or 1:50 CELLstart^TM^ (Thermofisher Scientific, A1014201)-coated tissue culture plastic (TCP) in a 48-well plate.

### 2.9 Cell Viability

After 48 hours of culture in proliferation medium on 2D substrates, mNSC viability was assessed using an ApoLive-Glo™ Multiplex Assay (Promega, G6410). This assay, which includes measurements of viability (ATP content) and apoptosis (caspase 3/7 activity), was run as per the manufacturer’s instructions. As a negative control for viability, mNSCs were seeded onto TCP and incubated with 10% ETOH for 10 minutes at 37°C to induce cell death immediately before absorbance measurements were acquired. A 48-well plate with all 2D materials, but no cells, was run in parallel to account for any background signal from either the luminogenic or fluorogenic substrates. In addition, cell viability was assessed using a LIVE/DEAD™ Viability/Cytotoxicity Kit (Invitrogen™, L3224), as per the manufacturer’s instructions. Images were acquired using a Zeiss Axio Observer 7 widefield fluorescence scope (10x air, 0.55 NA objective).

### 2.10 Immunocytochemistry and Image Analysis

mNSCs were cultured on 2D substrates, as described in 2.7, except that differentiation medium was used. On the fifth day of culture, half of the differentiation medium from each well was replaced with 4% paraformaldehyde to fix mNSCs to the substrates (15 minutes at room temperature). For immunocytochemistry, fixed mNSCs were permeabilized (0.1% v/v PBS-Triton, 5 minutes) and blocked (blocking solution – 2% bovine serum albumin, BSA, and 4% v/v normal donkey serum in 0.05% v/v PBS-Tween) for 1 hour at room temperature. Primary and secondary antibody solutions were diluted in the blocking solution (Supplementary Table 1). Samples were incubated with primary antibody overnight at 4°C. The following day, primary antibody was removed, and samples were rinsed and incubated with secondary antibody solution, including Hoescht (ThermoFisher Scientific, A-78952, 1:1000), for 1 hour at room temperature. Coverslips with fixed cells were mounted onto microscope slides (Globe Scientific, 1358W) using Fluoromount-G™ Aqueous Mounting Medium (Millipore Sigma, F4680). Images were acquired using a Zeiss Axio Observer 7 and a 20x air, 0.55 NA objective. Negative controls, which did not receive primary antibody, were also imaged. Image analysis was done using the open-source software CellProfiler. For each protein marker, raw grayscale images were thresholded at the same level to create a binary image. For nuclear markers (e.g., Ki-67, NeuN and Olig2), a watershed function was applied to thresholded images prior to counting the number of objects (i.e., labeled nuclei) in each image. For protein markers with non-nuclear expression (e.g., b-III-tubulin and GFAP), the total area of positive staining was calculated per image and divided by the total number of cells. Immunostaining quantification is reported here as either number of positively stained nuclei divided by total number of nuclei (Hoescht) or area of positive staining divided by total number of nuclei. Data is generated from 3-4 images per sample.

### 2.11 Statistical Analyses

GraphPad Prism 10 was used to create most data plots and conduct statistical analyses. The exception is for the water contact angle analysis, which was performed using MATLAB. Five WCA measurements were taken for each material, and a one-way ANOVA was performed between a) same material’s pre- and post-culture conditions, and b) post-culture material and post-culture bare-glass. For the ApoLive-Glo assay, background signal was subtracted from the mean fluorescence (living) and luminescence (dying) signal from mNSCs cultured on each material. These mean values were then normalized to readings from mNSCs cultured on PLO/laminin-coated coverslips. These values were then divided (living/dying) to give a measure of relative viability between the conditions. The standard error of the mean (SEM) was calculated across 4 independent repeats (each with n=4) and the error was propagated out accordingly. For the ApoLive-Glo data, the raw mean fluorescence values from mNSCs cultured on each material-coated coverslips were normalized to the raw mean fluorescence values from mNSCs cultured on PLO/laminin-coated coverslips. For all viability assessments, one-way ANOVA, followed by Tukey’s multiple comparison test was performed, to assess significant differences across experimental groups (Table S2). For data generated from immunocytochemistry images, one-way ANOVA followed by a Tukey’s multiple comparison test was also used (Table S2). A significance level (α) of 0.05 was used for all analyses.

## 3 RESULTS

### 3.1 Growth and Characterization of 2D Materials

To confirm the deposition of PtTe_2_, PtSe_2_, MoS_2_, and graphene onto glass substrates, structural characterization was performed using Raman spectroscopy. For graphene, two main characteristic peaks are observed, corresponding to the D-band at ∼1300 cm^−1^ and G-band at ∼1580 cm^−1^ (Fig. 2a)^31^. . PtSe_2_ samples exhibit two dominant peaks, with the E_g_ mode at 175 cm^-1^ and the A_1g_ mode at 205 cm^-1^, corresponding to the in-plane and out-of-plane vibration motions of the Se atoms, respectively (Fig. 2b)^29^. The Raman spectrum for CVD-grown PtTe_2_ shows peaks around 110 cm^-1^ and 157 cm^-1^, corresponding to E_g_ and A_1g_ modes, respectively (Fig. 2c)^28^Similarly, E^1^_2g_ (∼383 cm^−1^) and A_1g_ (408 cm^−1^) modes were observed where expected to confirm the presence of the MoS_2_ film or flaky MoS_2_ (Fig. 2d and Fig. 2e)^30^. To further validate these characterizations, we also measured Raman shifts for the bare-glass substrate (Fig. S6).

**Figure 2.**
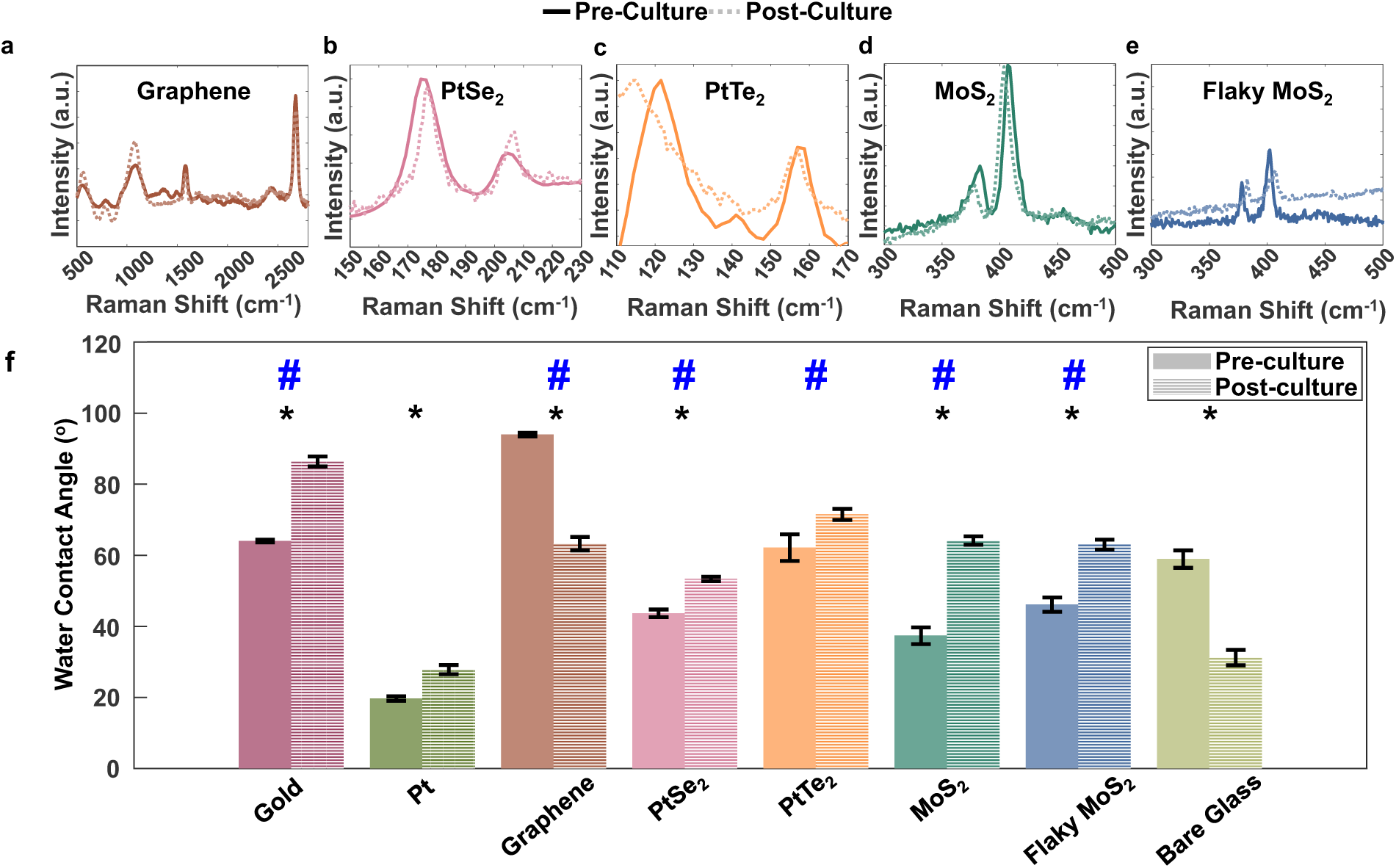
Raman spectra for a) Graphene, b) PtSe2, c) PtTe2, d) MoS2, and e) flaky MoS2. f) The water contact angle measurements for all materials both bare/pre-culture (solid) and post-culture (striped) (n = 5, mean +/- standard deviation). A considerable increase in water-contact-angles (WCA) after post cell-culture can be observed. The black * and blue # refer to statistical significance of difference in WCA between conditions (pre- and post-culture) and between a given material’s post-culture sample and the post-culture bare-glass substrate (p<0.0001), respectively.

One essential parameter of materials that are used for biological interfaces is their surface hydrophilicity/hydrophobicity, which can strongly influence protein adsorption and cell adhesion^32,33^. To characterize wettability of the 2D electronic materials, WCA measurements were performed on materials before and after protein coating and culture with cells (see Methods). After culture, as seen in Fig. 2f, post-culture, bare-glass had the lowest WCA of all the materials, except for Pt and flaky MoS_2_ (* p<0.0001). While graphene and bare glass showed considerable decreases in WCA post-culture, the rest of the materials had an increase in WCA (* p<0.0001). Further, with the exception of graphene, the materials in their pre-culture form had lower WCA values than post-culture, indicating that they were relatively hydrophilic prior to protein adsorption. Most of the freshly deposited electronic materials had a lower WCA than those on which cells had been previously cultured. Protein coatings are expected to decrease hydrophilicity as the hydrophobic material surface are most likely to interact directly with the proteins and the material would become hydrophobic due to the additional protein added upon the pristine material surface. The exceptions are graphene, which showed a considerably reduced WCA after cell culture, and PtTe_2_, which did not show a significant change in WCA both before and after cell-culture. These differences can likely be attributed to adsorption of proteins from serum in the culture medium to the material surfaces, which involves protein denaturation where the more hydrophobic regions of the protein associate with the hydrophobic regions of the material surface and the more hydrophilic regions of the protein associate with the aqueous culture medium. However, it is somewhat difficult to interpret the post-cell culture data given that trypsinization to detach the cultured cells likely also removes adsorbed proteins, likely to a different extent on different material surfaces.

### 3.2 Neural Stem Cell Biocompatibility

Viability and apoptosis of mNSCs seeded atop the 2D electronic materials were first assessed using an ApoLive-Glo Assay after 48 hours of culture in proliferation medium. Figure 3a shows the ratio of the live cell to apoptotic cell signal to give a measure of relative viability between the conditions. The relative-mean-fluorescence (representing the signal from living cells, proxy for ATP concentration) and -luminescence (representing the signal from apoptotic cells, proxy for caspase-3/7 activities) were normalized to readings from mNSCs cultured on PLO/laminin-coated glass coverslips. Figure 3b shows data from the same experiments but instead shows the fluorescence (live cell) signal from each group that was first normalized to a standard curve to calculate total number of living cells and then normalized to the number of living cells on the PLO/laminin-coated glass coverslip control. No statistically significant differences in the live/apoptotic cell ratio or the live cell signal were observed across the materials. This finding indicates that all the 2D electronic material substrates evaluated are at least as suitable as glass for mNSC culture.

**Figure 3.**
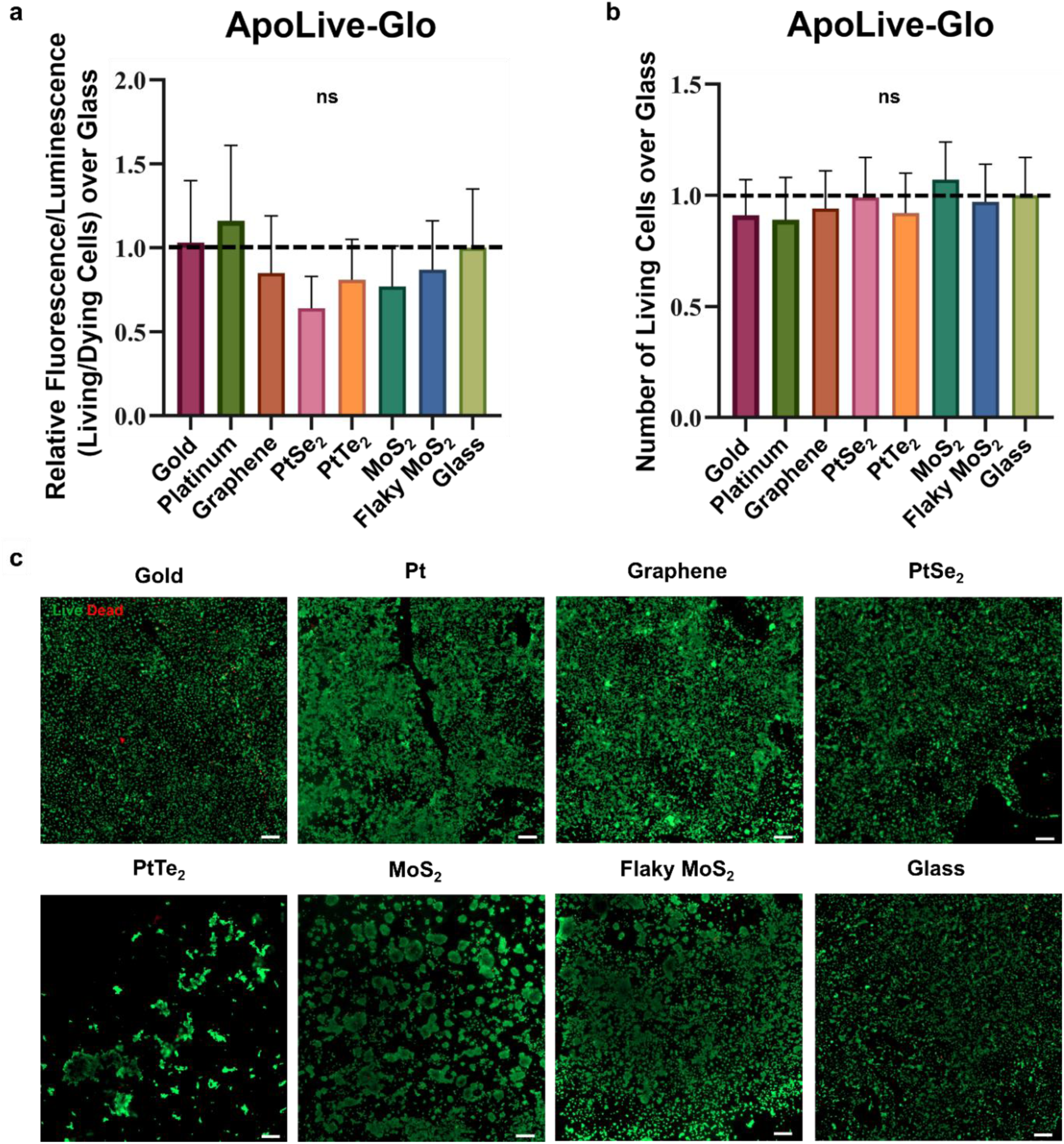
Viability of mNSCs after 48 hours of culture in proliferation medium. **a)** An ApoLive-Glo assay was performed to calculate percentage of relative fluorescence (living cells) over luminescence (dying cells) for each material. These values were normalized to the glass coverslip control condition and plotted (N=4 repeats in each of 4 independent repeats). Plot shows mean +/- SEM. **b)** The fluorescence values were also equated to the number of living cells based on a standard curve of serial dilutions within each repeat of the ApoLive-Glo assay. These values were normalized to the glass coverslip control condition and plotted (N=4 repeats in each of 4 independent repeats). Plot shows mean +/- SEM. One-way ANOVA showed no significant (ns) differences in viability when applied to data shown in (a) or (b). **c)** Representative images of the Live/Dead Assay showing live (Calcein AM, green) and dead (ethidium bromide, red) cells. Scale bars = 200 µm.

Further, a live/dead fluorescence imaging assay was performed for qualitative assessment of viability (Fig. 3c and Fig. S1). Visually, all groups showed similar proportions of living and dead cells. However, morphological differences in the appearance of the adhered cells were observed. While mNSCs formed monolayers on most substrates, including glass controls, they remained clustered on MoS_2_ and PtTe_2_ substrates. In general, poor mNSC adherence/spreading on PtTe_2_ substrates was consistently observed. Alternatively, flaky MoS_2_ showed relatively better adherence and more spreading of the cells across the material’s surface compared to its counterpart CVD-grown MoS_2._ PtSe_2_ showed relatively similar cell adherence to that of flaky MoS_2_ as well as the gold, platinum, graphene and glass control. Additionally, cells on the gold substrate appeared to spread less than other conditions while graphene appeared to promote the most clustering of live cells as evidenced by the very fluorescent green circular areas throughout the image.

To evaluate how each 2D electronic material may support differentiation, mNSCs were cultured for five days in differentiation medium. As mitogens are withdrawn with the introduction of differentiation medium, mNSCs are expected to stop dividing as they differentiate into neurons or glia. Ki-67, a nuclear protein expressed by cells in the G1, S, G2, or M phases of mitosis, was used to assess proliferative activity. mNSCs grown on flaky MoS_2_ had significantly more proliferative cells than those grown on glass coverslips (\**p* < 0.05) (Fig. S2). However, no conditions had more than around 12% of cells which were proliferative, indicating that many cells had exited the cell cycle to begin differentiation. Differentiation of mNSCs into neurons was assessed by expression of βIII- tubulin (neuronal precursors and mature neurons) (Fig. 4) and nuclear NeuN (mature neurons) (Fig. 5). There were no statistically significant differences in the expression of βIII-tubulin across the conditions (Fig. 4a). However, there was abundant expression of βIII-tubulin in all conditions, indicating that all the materials robustly support differentiation of mNSCs along the neuronal lineage at least as well as the glass control. Culture on flaky MoS_2_ substrates appeared to yield more mature neurons, as indicated by the presence of significantly more cells with nuclear NeuN expression, than PtTe_2_ (\*\**p* < 0.01), graphene (\**p* < 0.05), PtSe_2_ (\**p* < 0.05), and glass (\**p* < 0.05) substrates. While cultures on most material substrates had an average of 25-50% mature neuronal cells, the flaky MoS_2_ cultures had upwards of 75% neurons. While the mean percentage of mature neurons for CVD-grown MoS_2_ was lower than that for flaky MoS_2_, this difference was not statistically significant. It is possible that MoS_2_ itself promotes neuronal differentiation and this is further potentiated by the flaky conformation.

**Figure 4.**
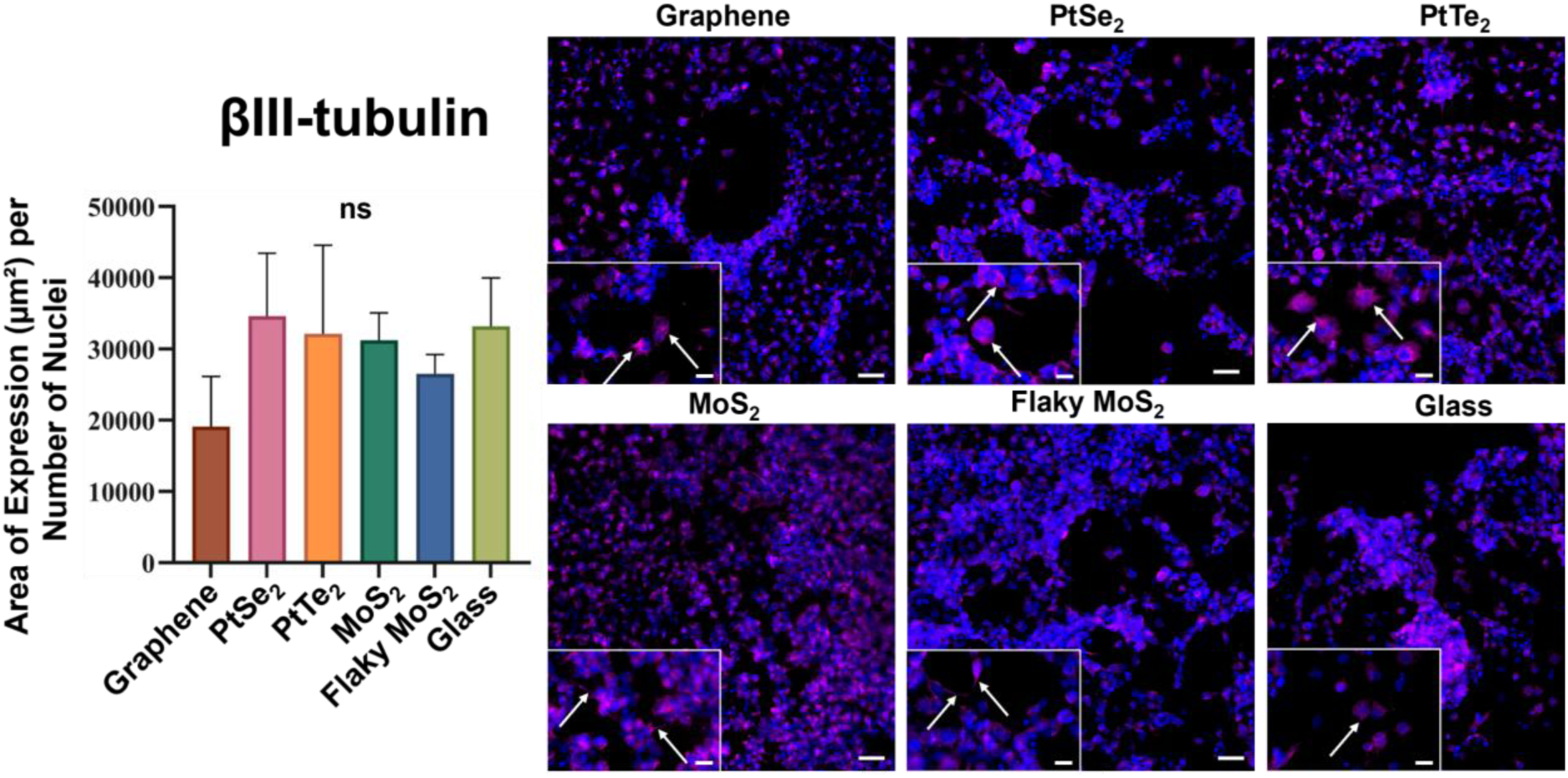
Differentiation of mNSCs towards neurons after 5 days in differentiation medium. βIII-tubulin (magenta) was used to identify both immature and mature neurons. Main scale bars = 50 μm and inset scale bars = 20 μm. Quantification shows βIII-tubulin+ area divided by total nuclei for each image (N=2 repeats in each of 2 independent repeats). One-way ANOVA showed no significant (ns) differences between the conditions. Plot shows mean +/- SEM.

**Figure 5.**
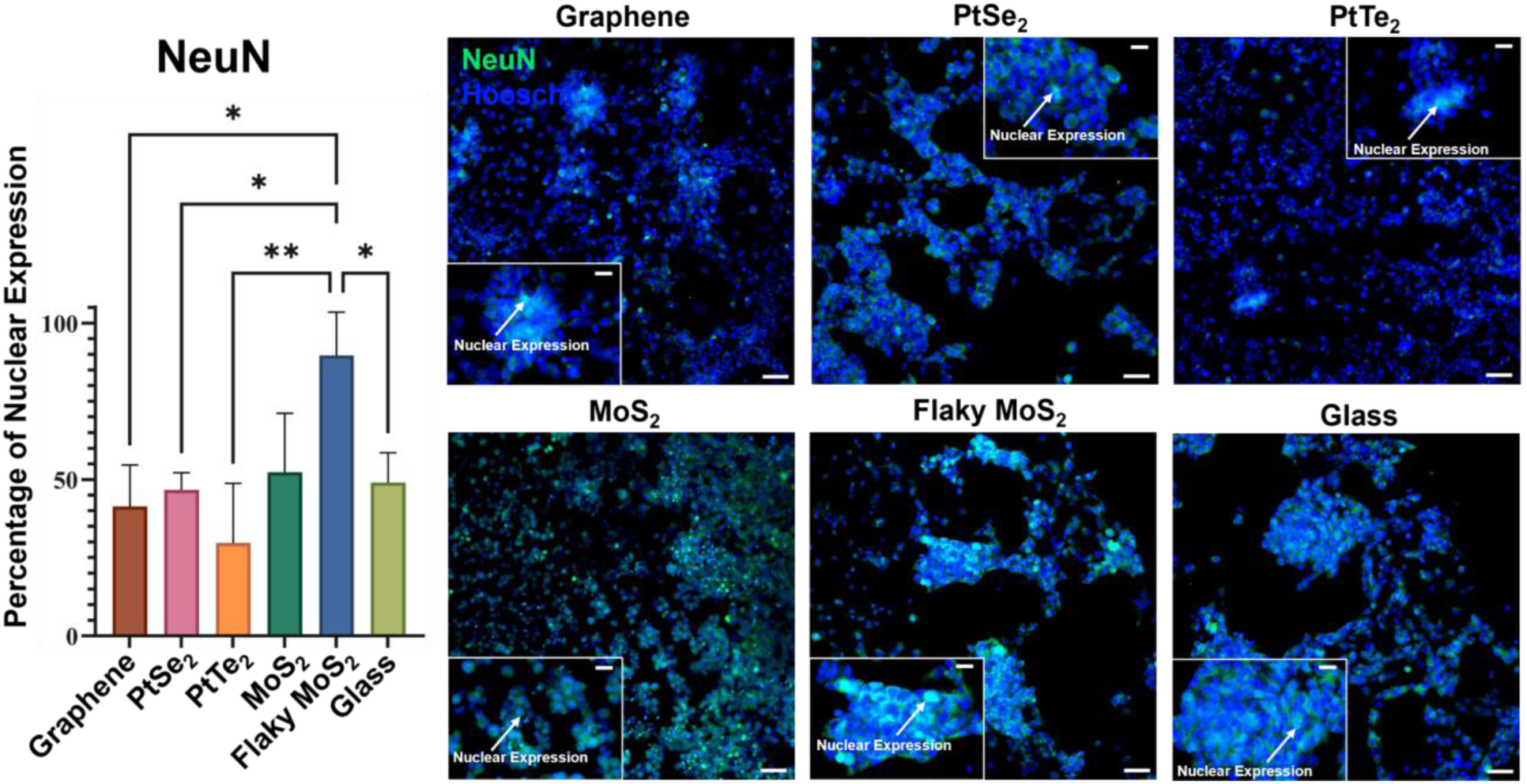
Differentiation of mNSCs into mature neurons after 5 days in differentiation medium. Nuclear expression of NeuN (green) was used to identify mature neurons. Main scale bars = 50 μm and inset scale bars = 20 μm. Quantification shows numbers of NeuN+ nuclei divided by total nuclei for each image (N=1,2 repeats in each of 2 independent repeats). One-way ANOVA showed that flaky MoS2 had significantly more nuclear expression than PtTe2 (\*\**p* < 0.01), Graphene (\**p* < 0.05), PtSe2 (\**p* < 0.05) and glass (\**p* < 0.05). All other conditions had no significant (ns) differences. Plot shows mean +/- SEM.

We further assessed the differentiation of mNSCs into mature glial cells. Interestingly, flaky MoS_2_ cultures had significantly fewer cells with nuclear expression of olig2, a marker of oligodendrocyte progenitors or early motor neuron progenitors, than did CVD-grown MoS_2_ (\*\**p* < 0.01) or PtSe_2_ substrates (\*\*\**p* < 0.001) (Fig. S3). PtSe_2_ cultures also had significantly more olig2-positive cells than cultures on graphene (\*\**p* < 0.01) or glass (\*\**p* < 0.01). PtSe_2_ and CVD-grown MoS_2_ substrates yielded statistically equivalent percentages of olig2-positive cells. Given the relatively low NeuN expression in the PtSe_2_ cultures (Fig. 5), it is more likely that the olig2-positive are oligodendrocytes, rather than motor neuron progenitors. For the CVD-grown MoS_2_ cultures, it is equally likely that the olig2-positive cells represent oligodendrocytes or motor neuron progenitors. Given the low numbers of olig2-positive cells relative to the numbers of NeuN-positive cells on flaky MoS_2_ cultures, mNSCs likely differentiated towards neurons rather than oligodendrocytes. For these specific assays, PtTe_2_ did not show enough adherence and hence had to be excluded from analysis.

Differentiation into astrocytes was assessed via expression of glial fibrillary acidic protein (GFAP) (Fig. S4). PtSe_2_ cultures had significantly more cells expressing GFAP than flaky MoS_2_ cultures (\**p* < 0.05). All other conditions showed similar levels of GFAP expression. Together, these results indicate that the PtSe_2_ material promotes preferential differentiation of glia, namely oligodendrocytes and astrocytes, whereas the flaky MoS_2_ material promotes neuronal differentiation.

## 4 DISCUSSION

Neural-bioelectronic interfaces are potentially transformative for *ex vivo* applications such as platforms for drug discovery and development as well as *in situ* applications such as implanted devices for therapeutic stimulation of the nervous system. A primary requirement of any electrically conductive bioelectronic material is its biocompatibility. Beyond preserving neural cell viability, effects of these materials on neural cell phenotype are crucial to characterize. Here, we evaluated the potential of several 2D electronic materials for interfacing with primary mNSCs.

Our findings demonstrate that large-area, CVD-grown 2D electronic materials including graphene, PtSe₂, PtTe₂ and MoS₂, exhibit biocompatibility with mNSCs, supporting both cell viability and differentiation without inducing cytotoxic effects. In contrast, several prior studies that primarily investigated biocompatibility of some these electronic materials as exfoliated flakes, as opposed to the CVD-grown 2D format, have reported variable results, perhaps depending on material type, degree of exfoliation, and cell type tested (Table S3). For example, exfoliated MoS_2_ and WS_2_ have been shown to damage plasma membranes, disrupt cytoskeletal integrity, and induce inflammatory responses in non-neuronal, transformed cell lines, including the A549 lung carcinoma line, the THP-1 human monocyte line, and the RAW264.7 mouse macrophage lines^24,45,58^. Similarly, greater exfoliation of MoS_2_ nanosheets has been correlated with increased toxicity^53^, in theory because exfoliated nanoparticles or flakes are more cytotoxic when engulfed by cells^60^. In neural contexts, some reports have indicated beneficial effects of MoS_2_ flakes on rat NSC attachment without measurable toxicity^43^, but these studies lacked direct comparisons across multiple 2D materials or large-area formats. Our work expands upon previous findings by showing that material format (flaky vs. CVD-grown) significantly impacts bioactivity, as flaky MoS_2_ enhanced neuronal differentiation while CVD-grown MoS₂ did not elicit the same effect. MoS₂ flakes have gaps between them where the underlying glass is exposed. Since the MoS₂ flakes were applied via spin-coating, cells may encounter flat, flake-free regions of glass that promote adhesion and survival, potentially contributing to the higher observed viability and neuronal differentiation. While previous studies have largely confirmed that graphene can support neuronal viability and neurite outgrowth^48–51^, our comparative analysis highlights that several other 2D materials can similarly support neural cell culture. To our knowledge, PtSe₂ and PtTe₂ have not previously been explored in biocompatibility studies involving cultured stem cells or neurons, making our work the first to evaluate their effects on neural differentiation. Furthermore, we were able to identify materials, like PtSe_2_, that preferentially affect lineage specification. Collectively, these results suggest that large-area 2D materials may offer superior and more predictable biocompatibility for neural interfacing applications than their exfoliated counterparts, underscoring the importance of material format and substrate continuity in bioelectronic design.

It is important to understand how a bioelectronic interface may change or degrade during cell-culture as this may impact its mechanical properties or lead to the release of cytotoxic by-products. We first used Raman spectroscopy to validate whether the materials degraded/ washed-off after undergoing cell-culture and assays and found all the materials of interest to be intact over the substrates. We also performed WCA measurements to check surface wettability both pre- and post-culture. We saw an increase in WCA across all materials post-culture, except graphene which decreased, and PtTe_2_ which had no significant changes in WCA between pre- and post-cultures. The considerably lower WCA across all materials compared to the bare-glass substrate implies that the surface is conducive for protein coatings and cell adherence. The consistent increase in WCA after cell-culture might potentially indicate the presence of hydrophobic protein-cores pointing away from the material surface. Since the adhesion protein that best promotes cell adhesion may vary for different 2D materials and the cell-type being tested, a future direction of this research could be a longitudinal study where various proteins (e.g., poly-L-lysine, poly-L-ornithine, fibronectin, collagen or laminin) are applied at various concentrations to fundamentally study WCAs for each material.

2D materials have a smoother surface compared to their bulkier counterparts owing to their atomically thin single-layer to few-layers. Cells usually prefer rougher surfaces to adhere and proliferate^34,35^. While surface roughness was not quantified in this study, the flaky MoS_2_ substrates are likely to be rougher than the other substrates because of the method by which the MoS_2_ flakes were adhered together on top of a smoother glass substrate. This increased roughness might explain improved cell adhesion and increased cell proliferation in the flaky MoS_2_ compared to the CVD-grown MoS_2_ cultures. This hypothesis could be further investigated in future studies using atomic force microscopy to visualize the cell adherence patterns across the surface of the material. However, above a certain roughness or quantity of surface-defects, the material can potentially become cytotoxic through mechanical damage to cells or through being internalized by the cells^23–25^. This does not appear to be the case with the flaky MoS_2_ substrates given the comparable viability of mNSCs on all substrates evaluated. While the flaky MoS_2_ substrate was the only condition on which mNSC proliferation was higher than the glass control, it also supported the most neuronal differentiation. This may seem counterintuitive, as stem cells typically stop proliferating as the differentiate. It is possible that a smaller subpopulation of proliferative mNSCs remains in these cultures alongside a larger mature neuronal population. Importantly, these results show that the method by which MoS_2_ is deposited (i.e., flaky vs CVD-grown) strongly influences the materials’ bioactivity. Few glial cells were observed on the flaky MoS_2_ substrate. In contrast, the PtSe_2_ substrate appeared to promote oligodendrocyte-lineage and astrocyte-lineage differentiation.

Taken together, for future bioelectronic interfaces, graphene and MoS_2_ are promising scaffolds for applications requiring a mixed population of immature neuronal and mature glial cells. Owing to their electronic characteristics, they can potentially be used as electrode materials for *in vitro* platforms and *in vivo* stimulating/recording electrodes. Further, MoS_2_ and flaky MoS_2_ are best suited for applications that require more mature neuronal cultures such as *in vitro* studies of neural network characterizations. In the future, more 2D materials in their CVD-grown form should be characterized for their biocompatibility and electrophysiological recording capabilities, to create a library of such materials for bioelectronic interface applications.

## 5 CONCLUSION

This study characterized mNSC viability and differentiation potential when cultured on an array of CVD-grown or flaky, electronics-grade 2D materials that included important candidates for bioelectronic and neural interfacing (i.e., graphene, PtSe_2_, PtTe_2_, MoS_2_, flaky MoS_2_). All bioelectronic materials supported viability of cultured mNSCs as well as conventional culture substrates, namely laminin-coated glass. We found that the material substrate strongly influenced mNSC differentiation, where flaky MoS_2_ substrates induced neuronal maturation while PtSe_2_ substrates promoted maturation of glial cells, including oligodendrocyte progenitors and astrocytes. Strikingly, the format (i.e., flaky vs CVD-grown) of MoS_2_ appeared to influence neuronal differentiation more than the chemistry, as only flaky MoS_2_ preferentially promoted neuronal maturation over other material types. In contrast to previous reports, this study provides a comprehensive comparison of promising candidate materials for bioelectronic interfacing in large-area, 2D formats.

## ACKNOWLEDGMENT

We would like to acknowledge all the members of the Akinwande Nano Research Lab, and Seidlits Lab, in providing their valuable suggestions and feedback. BioRender was used to create aspects of some figures.

