## Supplementary Materials for "BIOCOMPATIBILITY OF LARGE-AREA 2-DIMENSIONAL ELECTRONIC MATERIALS WITH NEURAL STEM CELLS"

### SUPPLEMENTARY SECTION

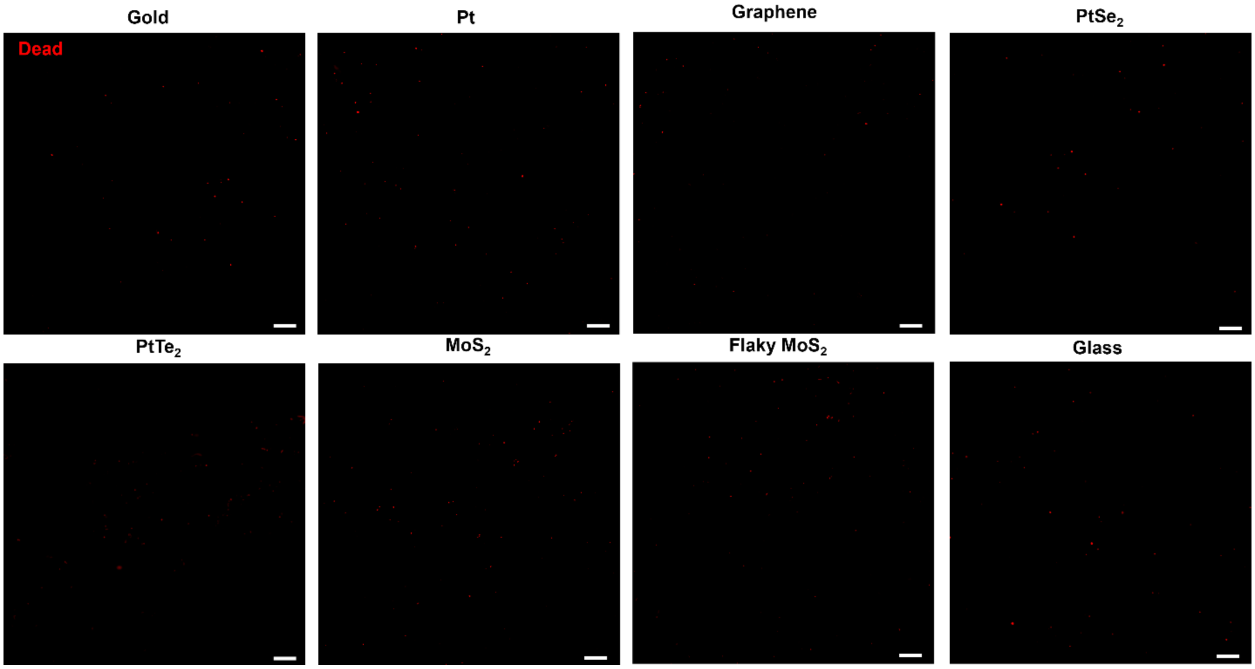


**Figure S1.** Viability of mNSCs after 48 hours of culture in proliferation medium. Representative images of the Live/Dead Assay showing just the dead (ethidium bromide, red) cells. Scale bars = 200 μm.

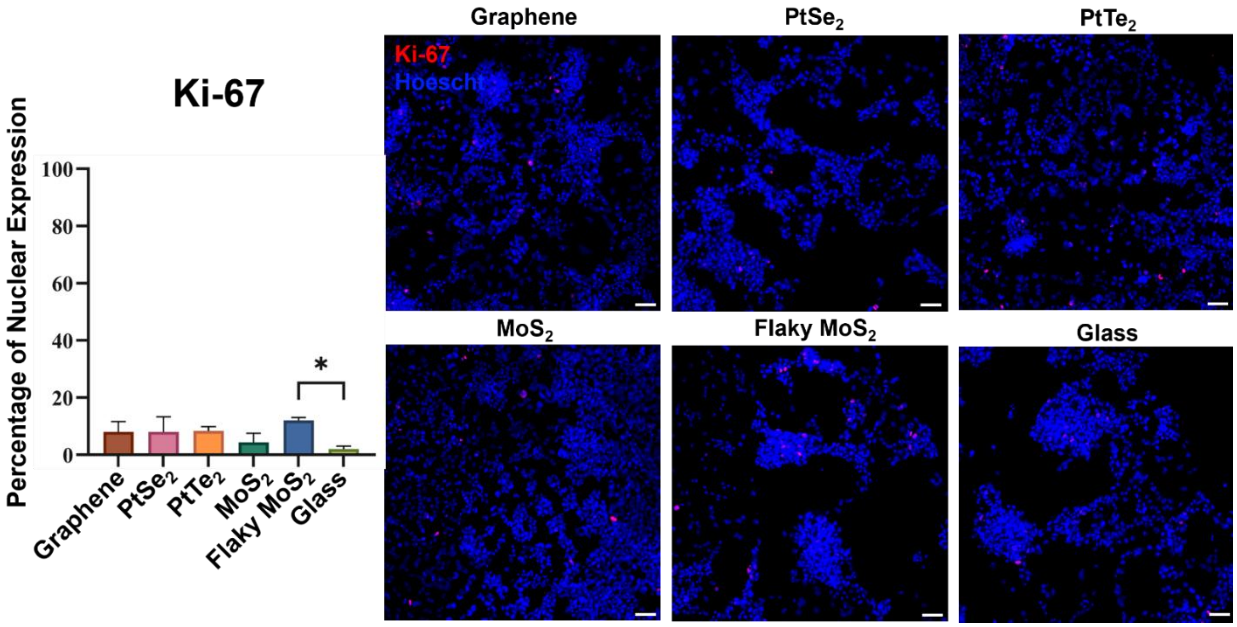


**Figure S2.** Proliferation of mNSCs after 5 days in differentiation medium. Ki-67 (red) staining was performed to evaluate the proportion of proliferative cells for each condition. Scale bars = 50. Quantification shows Ki-67+ area divided by total nuclei for each image (N=1,2 repeats in each of 2 independent repeats). One-way ANOVA shows mNSCs grown on flaky $MoS_2$ had significantly more nuclear expression of Ki-67 than those grown on the glass control condition (*$p < 0.05$). All other conditions had no significant (ns) differences. Plot shows mean +/- SEM.

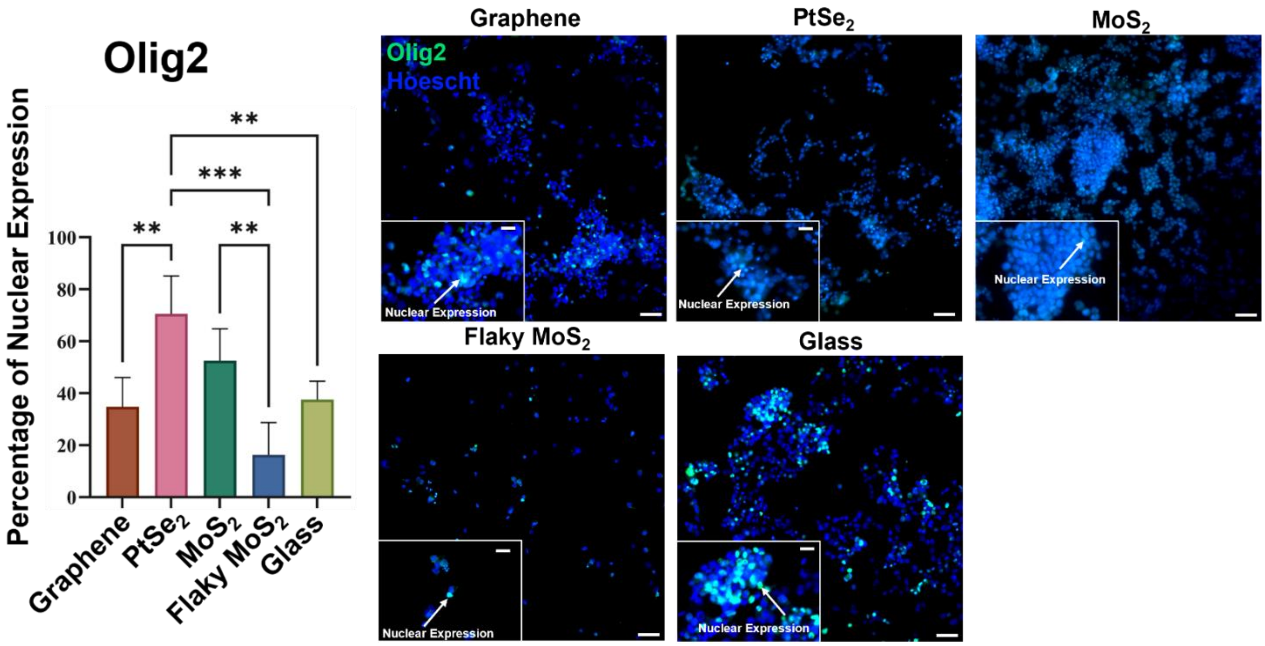


**Figure S3.** Differentiation of mNSCs towards oligodendrocyte or motor neuron precursors after 5 days in differentiation medium. Olig2 (green) was used to identify cells that were either oligodendrocyte or motor neuron precursors. Main scale bars = 50 µm and inset scale bars = 20 µm. Quantification shows Olig2+ area divided by total nuclei for each image (N=2 repeats in each of 2 independent repeats). One-way ANOVA showed that $PtSe_2$ had significantly more nuclear expression of Olig2 than graphene and glass (**$p < 0.01$), as well as flaky $MoS_2$ (***$p < 0.001$). One-way ANOVA also showed that $MoS_2$ showed significantly more nuclear expression of Olig2 than its counterpart flaky $MoS_2$ (**$p < 0.01$). All other conditions had no significant (ns) differences. Plot shows mean +/- SEM.

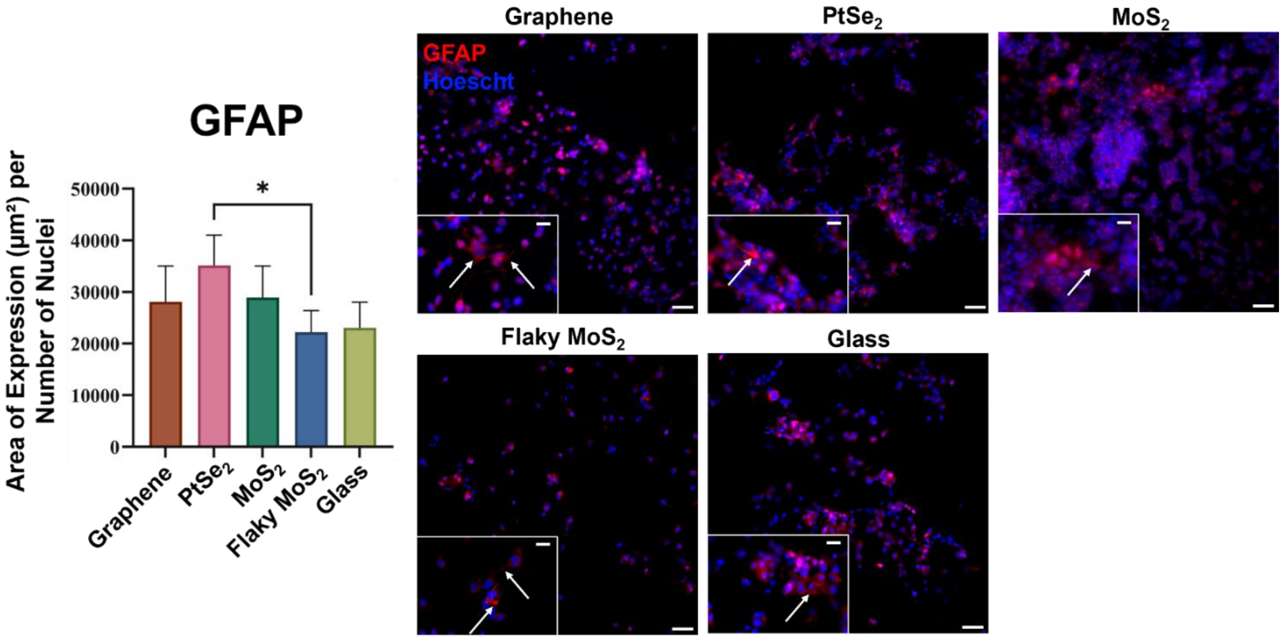


**Figure S4.** Differentiation of mNSCs towards astrocytes after 5 days in differentiation medium. GFAP (red) was used to identify cells that were either astrocytes. Main scale bars = 50 µm and inset scale bars = 20 µm. Quantification shows GFAP+ area divided by total nuclei for each image (N=2 repeats in each of 2 independent repeats). One-way ANOVA showed that $PtSe_2$ showed significantly more GFAP expression per number of nuclei compared to flaky $MoS_2$ (*$p$ < 0.05). All other conditions had no significant (ns) differences.

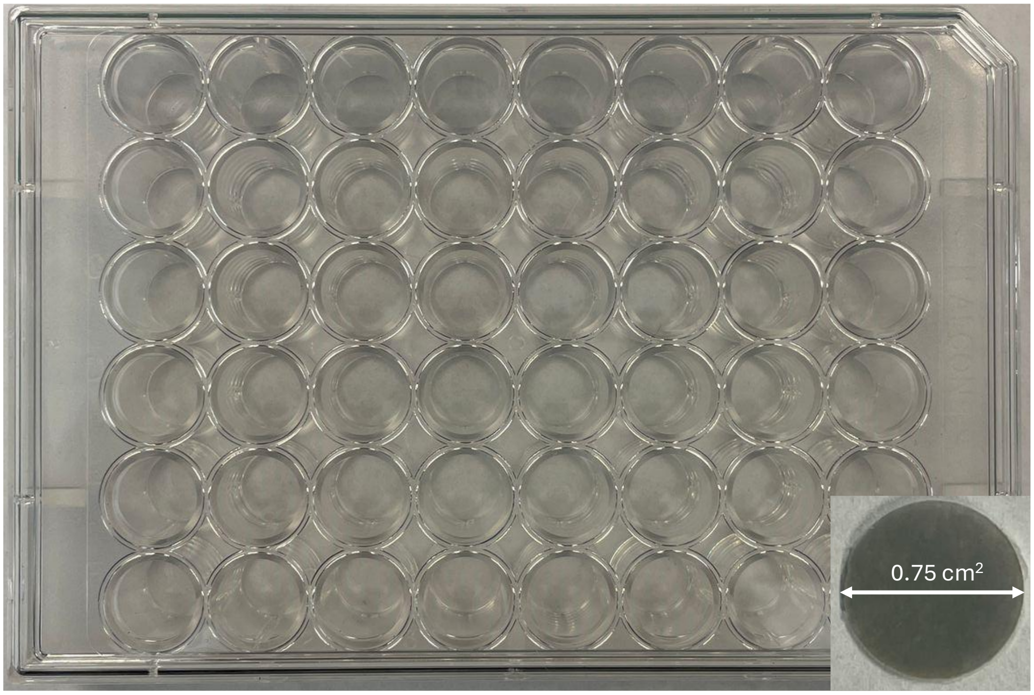


**Figure S5.** A 48-well plate that was used for all the cell-viability assays in the present study. 4 sample of each material-type were placed at the bottom of the well-plate. Inset: And example glass slip containing the 2D material of interest (in this case, $PtSe_2$)

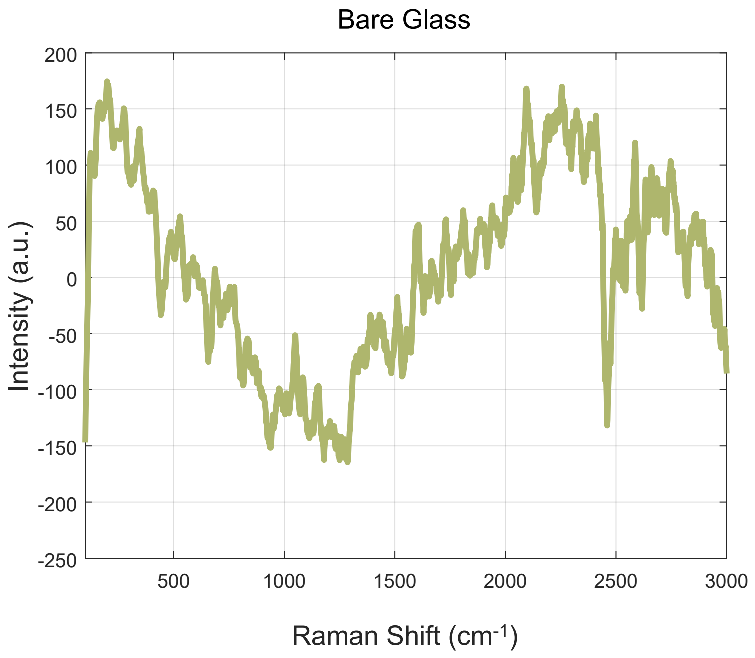


**Figure S6.** Raman Spectrum of the bare glass substrate.

**Table S1.** Immunocytochemistry antibodies and dilutions.

| Antibody | Manufacturer | Catalog Number | Target | Dilution |
|---|---|---|---|---|
| Ki67 | Novus | NB500-170 | Proliferative cells | 1:100 |
| NeuN | Millipore Sigma | MAB377 | Neuronal cells | 1:100 |
| B-III-Tubulin | Millipore Sigma | AB9354 | Early neuronal cells | 1:1000 |
| Olig2 | Millipore Sigma | MABN50 | Oligodendrocytic cells | 1:500 |
| GFAP | Novus | NB300-141 | Astrocytic cells | 1:1000 |

**Table S2:** One-way ANOVA values, adjusted p values based on Tukey's post-hoc analyses, for viability assays and image quantification performed in this research study.

| Measurement | Adjusted P Value | Figure |
|---|---|---|
| **ApoLive-Glo:** Relative Fluorescence/Luminescence (Living/Dying Cells) over Glass | All conditions had no significant (ns) differences | Figure 3a |
| **ApoLive-Glo:** Number of Living Cells over Glass | All conditions had no significant (ns) differences | Figure 3b |
| **βIII-tubulin** | All conditions had no significant (ns) differences | Figure 4 |
| **NeuN** | Flaky $MoS_2$ > $PtTe_2$ (**$p < 0.01$), Flaky $MoS_2$ > Graphene (*$p < 0.05$), Flaky $MoS_2$ > $PtSe_2$ (*$p < 0.05$), Flaky $MoS_2$ > glass (*$p < 0.05$), All other conditions had no significant (ns) differences | Figure 5 |
| **Olig2** | $PtSe_2$ > graphene (**$p < 0.01$), $PtSe_2$ > glass (**$p < 0.01$), $PtSe_2$ > flaky $MoS_2$ (***$p < 0.001$). $MoS_2$ > flaky $MoS_2$ (**$p < 0.01$), All other conditions had no significant (ns) differences | Figure S3 |
| **GFAP** | $PtSe_2$ > flaky $MoS_2$ (*$p < 0.05$), All other conditions had no significant (ns) differences. | Figure S4 |
| **Ki-67** | Flaky $MoS_2$ > glass (*$p < 0.05$), All other conditions had no significant (ns) differences | Figure S2 |

**Table S3.** An exhaustive table of previous biocompatibility studies with 2D materials, in comparison with the present study.

| Ref | 2D Material (Flaky and CVD-grown) | Cell Type | Cytotoxicity Test | Conclusion |
|---|---|---|---|---|
| **This work** ★ | CVD-grown 2D Electronic materials – Graphene (G), $MoS_2$, $PtSe_2$, $PtTe_2$, metals – Gold, Pt, non-CVD 2D material – flaky $MoS_2$ | Mouse Neural Stem Cells | Apo-Live Glo assay, Fluorescence live-dead imaging, Immunocytochemistry for Astrocytes and Oligodendrocytes | All CVD-grown 2D electronic materials show excellent biocompatibility and are conducive to neural stem cell differentiation |
| **23** | Exfoliated $MoS_2$, $WS_2$, $WSe_2$, and G | Human lung Carcinom a Epithelial cells | MTT and CCK8 assays | $WSe_2$ showed similar degrees of cytotoxicity; GO and halogenated G are more toxic |
| **24** | Exfoliated $WS_2$ nanosheets | RAW264. 7 and A549 cell lines | CCK8 assay, LDH assay, and immunocytochemistry | $WS_2$ can severely damage plasma membrane and cytoskeleton |
| **25** | Exfoliated $VS_2$ , $VSe_2$ and $VTe_2$ | Human lung carcinom a (A549) cells | MTT and WST-8 assays | $VS_2$ , $VSe_2$ and $VTe_2$ are more toxic compared to $MoS_2$ |
| **45** | $MoS_2$ flakes | THP-1, A549, and AGS cells | Fluorescence Microscopy | $MoS_2$ caused Inflammatory responses in the cells |
| **53** | $MoS_2$ flakes | Human lung carcinom a epithelial cell line A549 | MTT and WST-8 assays | Higher exfoliation results in higher cytotoxicity |
| **54** | GO, hexagonal-Boron Nitride, $MoS_2$ | Primary human dendritic cells | Cell surface markers such as CD80, CD86, CD83, and CD40 | GO, hBN and $MoS_2$ are cytotoxic compared with the standard tissue-culture plate control |
| **55** | $MoS_2$ nanosheets | None | None | Simulation results: Peptide structural distortions upon binding to $MoS_2$ nanosheets |
| **56** | Exfoliated $MoS_2$ nanosheets | Mast cells | Electron microscopy and degranulation assay | $MoS_2$ is biocompatible |
| **57** | G, GO, $MoS_2$, and $WS_2$ | Intestinal epithelial cells | Multi-angle laser diffraction, XPS | Digestion/internalization of flakes observed |
| **58** | Exfoliated $MoS_2$ and BN flakes | Human Hepatom a HepG2 cells | Mitochondrial depolarization, and membrane integrity | $MoS_2$ and BN damage plasma membrane and increase arsenic toxicity |
| **59** | Exfoliated MoS2 and BN flakes | HDFn cells | LDH assay | Significant mitochondrial damage by $MoS_2$ and BN |
| **43** | $MoS_2$ Flakes | Rat Neural Stem Cells (NSC) | Fluorescence Microscopy | $MoS_2$ shows positive effect on rNSC attachment without measurable toxicity |

| | | | | |
|---|---|---|---|---|
| **44** | Exfoliated $WS_2$, $MoS_2$, CVD-grown $MoS_2$ | HEK293f cells | Fluorescence Microscopy | The materials are not deleterious to cellular viability or induce genetic defects |
| **46** | $MoS_2$ flakes | PC12 | SRB Assay | $MoS_2$ nanosheets are biocompatible |
| **Graphene** | | | | |
| **38** | G | Cos-7 cells, Primary hippocampal neurons | Fluorescence microscopy and stains. | G accelerates MSCs outgrowth |
| **39** | G, GO | Bone marrow derived MSCs | Immunofluorescence Imaging | G and GO promote cell adhesion and proliferation with no detectable cell stress |
| **41** | G, GO | Mouse iPSCSs 20D17 with GFP | Hemocytometry, qRT-PCR, and gene transfer analysis | G, GO yields spontaneous differentiation of iPSCs without significant disparity |
| **42** | G and Multi-walled Nanotube (MWNT) | H9 hESCs | Fluorescence Microscopy | The hESCs remained viable and pluripotent on G and MWNT–G hybrid |
| **47** | Single Layer G (CVD-grown) | Primary cortical neurons | Immunofluorescence Imaging | Both SLG and patterned SLG preserve synaptic efficacy |
| **48** | G | Primary rat cortical neurons | LDH Assay, Fluorescence and Brightfield Microscopy | Graphene is not more cytotoxic than the bare control surface |
| **49** | G, Carbon Nanotube (CNT) | Mouse Hippocampal neurons | SEM and Fluorescence Microscopy | G is a permissive interface, even when uncoated by cell adhesion layers, retaining unaltered neuronal signaling properties |
| **50** | G + Poly-L-Lysine, and bare G | Mouse hippocampal neurons | Fluorescence Microscopy | Crystalline quality of G tunes the neuronal affinity |
| **51** | G | Mouse hippocampal neurons | Immunofluorescence Imaging | G boosts neurite sprouting and outgrowth |
